# A collection of genetic mouse lines and related tools for inducible and reversible intersectional misexpression

**DOI:** 10.1101/754192

**Authors:** Elham Ahmadzadeh, N. Sumru Bayin, Xinli Qu, Aditi Singh, Linda Madisen, Daniel Stephen, Hongkui Zeng, Alexandra L. Joyner, Alberto Rosello-Diez

## Abstract

Thanks to many advances in genetic manipulation, mouse models have become very powerful in their ability to interrogate biological processes. In order to precisely target expression of a gene of interest to particular cell types, intersectional genetic approaches utilizing two promoter/enhancers unique to a cell type are ideal. Within these methodologies, variants that add temporal control of gene expression are the most powerful. We describe the development, validation and application of an intersectional approach that involves three transgenes, requiring the intersection of two promoter/enhancers to target gene expression to precise cell types. Furthermore, the approach utilizes available lines expressing tTA/rTA to control timing of gene expression based on whether doxycycline is absent or present, respectively. We also show that the approach can be extended to other animal models, using chicken embryos. We generated three mouse lines targeted at the *Tigre* (*Igs7*) locus with TRE-loxP-tdTomato-loxP upstream of three genes (*p21, DTA* and *Ctgf*) and combined them with Cre and tTA/rtTA lines that target expression to the cerebellum and limbs. Our tools will facilitate unraveling biological questions in multiple fields and organisms.

**Summary statement:** Ahmadzadeh et al. present a collection of four mouse lines and genetic tools for misexpression-mediated manipulation of cellular activity with high spatiotemporal control, in a reversible manner.

## INTRODUCTION

The wealth of genetic tools that have been developed for mouse to study biological questions *in vivo* have enabled extremely sophisticated experiments to be conducted. Especially useful are tools that have a modular nature. One approach for inducing gene mis-expression is to combine a mouse line carrying a driver module that expresses a protein (transcription factor) that induces gene expression only when the right genetic elements (DNA binding sites) are present with a line carrying an effector module that contains the necessary genetic elements upstream of a gene of interest (GOI). In this way only the offspring combining both modules will specifically express the GOI in particular cells defined by the driver module (Kaczmarczyk and Jackson, 2015; Rosen et al., 2015). This system can be modified to implement temporal and spatial control of gene expression by making it dependent on the activity of site specific recombinases (SSRs) (e.g. Cre), and by expressing SSRs from gene promoters or loci with specific expression patterns (Smedley et al., 2011). One drawback of using SSR systems, including inducible versions that allow for temporal control (e.g. CreER) (Brocard et al., 1998; Feil et al., 1997; Feil et al., 2009; Metzger et al., 1995), is that recombination of the target sites (loxP in the case of Cre) is irreversible. Hence, if misexpression of a GOI is dependent only on Cre activity, it cannot be reversed, and therefore studying any recovery process after a transient genetic manipulation is not possible.

The possibility of reversibility has been resolved by using other driver-effector combinations, the most prevalent in mouse being the Tet system, based on the tetracycline (Tet) resistance operon of *E. coli* (Hillen and Berens, 1994). In the first version of this system, a transactivator fusion protein composed of the tetracycline repressor (tetR) and the activating domain of the herpes simplex viral protein 16 (VP16) regulate the expression of a GOI by placing a Tet-responsive element (TRE) containing tet-operator sequences (tetO) upstream of the gene sequence. This system is known as Tet-Off, because in the presence of tetracycline or its analog doxycycline (Dox), the affinity of tetR to *tetO* sequences diminishes by 9 orders of magnitude, so that expression of the gene of interest is greatly reduced (Gossen and Bujard, 1992; St-Onge et al., 1996). Conversely, the Tet-On system is based on a reverse tetracycline-controlled transactivator (*rtTA*) derived from tTA through four amino acid changes in the tetR DNA binding moiety. This alters the binding properties of rtTA such that it can only recognize the *tetO* sequences in the presence of Dox (Gossen et al., 1995; Urlinger et al., 2000). A problem that for many years encumbered systems using tTA and rtTA (referred to as (r)tTA) was that the TRE DNA sequences can be inactivated through epigenetic mechanisms, rendering them unreliable unless de-repressed by additional genetic modules (Godecke et al., 2017). This drawback was overcome when a systematic screen identified a locus in the mouse genome that does not show this inactivation (Zeng et al., 2008). Indeed, *Tigre* (aka *Igs7*) is an intergenic region on mouse chromosome 9 that provides a safe harbor from which TRE-dependent expression can be <u>tig</u>htly <u>re</u>gulated (hence the name).

Despite these advances, all systems are limited by the fact that individual promoters/enhancers or gene loci often drive the expression of Cre/(r)tTA in a broader area (or more cell types) than what is required for the experiment, leading to insufficient specificity. Several intersectional approaches have been developed that allow for greater control over what cells are genetically manipulated. In one approach separate fragments of the Cre recombinase (known as “split cre”) are expressed from distinct drivers (promoter/enhancer sequences), resulting in Cre activity (loxP excision) only in the intersectional cell population where both halves of Cre are expressed (Hirrlinger et al., 2009). Another intersectional approach utilizes responder modules that depend on two types of recombinases (e.g. Cre and Flp), such as the systems developed by the Dymecki lab (Dymecki and Kim, 2007). However, neither approach allows for the GOI expression to be regulated in a reversible manner.

One strategy that allows for both precision of cell-type targeting and reversibility of gene mis-expression was recently developed by our labs (Madisen et al., 2015; Rosello-Diez et al., 2018). The approach involves combining the Cre/loxP and Tet systems such that the *Tigre* locus contains a double conditional allele in which expression of the GOI depends both on previous exposure to Cre (to prime the system through elimination of a floxed STOP sequence upstream of the GOI) and on the activity of (r)tTA (through a TRE sequence upstream of the loxP-STOP-loxP-GOI-polyA cassette). The version of this system that includes a tdTomato (tdT) coding sequence for the floxed-STOP cassette was called DRAGON (Fig. 1A), standing for Doxycycline-dependent and Recombinase-Activated Gene OverexpressioN (Rosello-Diez et al., 2018). The GOI in the first DRAGON line was the cell-cycle suppressor gene *p21* (*Tigre*^*Dragon-p21*^) and it was used to study cellular compensation mechanisms in the developing limb bones upon cell cycle blockade (Rosello-Diez et al., 2018). Since the DRAGON approach relies on two promoter/enhancers, it is intersectional (Fig. 1B), enabling the study of specific tissue-type dependent effects of gene misexpression. In addition, due to its dependence on (r)tTA, the system is inducible and reversible (Fig. 1C, D), allowing the effect of different times and durations of misexpression to be tested. Finally, by developing additional modules with particular GOIs involved in a specific cellular process (e.g. to reduce cell numbers by either blocking proliferation or inducing cell death), the system can effectively determine how different types of insults influence a biological process. Here we describe a series of new *Tigre*^*Dragon*^ lines (Fig. 1E) and provide a proof-of-principle for the kind of uses that they enable in two developing organs, the cartilage that drives long-bone growth in the limb (Kronenberg, 2003) and the excitatory cells of the cerebellar nuclei required for cerebellum growth (Willett et al., 2019). We also describe non-intersectional applications for comparison purposes, and the possibility of using these tools for transient gene misexpression in avian models.

**Figure 1.**
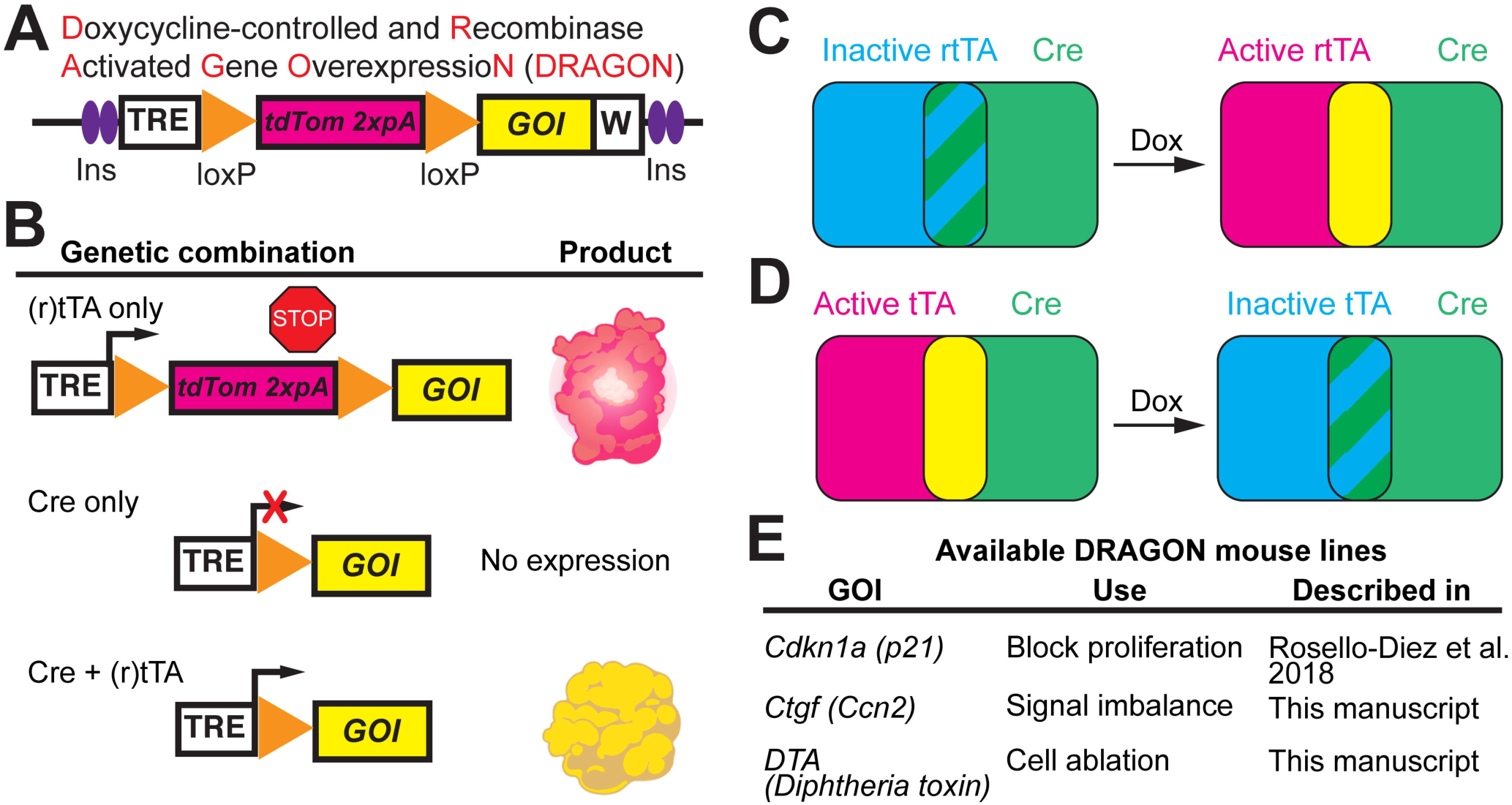
An intersectional genetic system for inducible and reversible gene misexpression. A) Basic components of the *Tigre*^*Dragon*^ alleles. Ins: insulators. W: Woodchuck virus RNA stabilizing sequence (WPRE). Expression of the Gene of interest (GOI) depends on both Cre and (r)tTA. B-D) In the presence of just (r)tTA (blue color in C and D), only the tdTomato (tdT) embedded in the intact floxed cassette is expressed (top row in B, magenta color in C and D). In the presence of just Cre (green color in C and D), the floxed tdTomato-STOP cassette is eliminated, priming the system and deleting tdT, but the GOI is not expressed (middle row in B). It is only in the presence of both Cre and (r)tTA (hatched blue and green in C and D) that the GOI is expressed (bottom row in B, yellow color in C and D). As the expression of both tdT and the GOI depends on (r)tTA activity, their expression is inducible and reversible, as it requires the presence of Doxycycline (Dox) for rtTA to be active (C), or its absence in the case of tTA (D). E) Table summarizing the available *Dragon* lines and their potential uses.

## RESULTS

### A *Tigre*^*Dragon-DTA*^ model can be used to generate acute tissue-specific cell ablation

A common theme in developmental and regenerative biology is that different responses are elicited by cell ablation compared to defective cell proliferation. Therefore, to complement our *Tigre*^*Dragon-p21*^ line, we generated an intersectional *Tigre*^*Dragon-DTA*^ model (see Methods, Fig. 2A and (Willett et al., 2019)) to enable cell ablation with tissue specificity and temporal control, via expression of attenuated <u>d</u>iphtheria toxin fragment <u>A</u> (Breitman et al., 1990). To kill some of the excitatory cerebellar nuclei neurons (eCN) that are the main output neurons of the cerebellum, we combined it with two transgenes: *Atoh1-tTA* (Tet-Off) (Willett et al., 2019) and *Atoh1-Cre* (Matei et al., 2005). Both transgenes are expressed in cells after they leave the progenitor zone called the rhombic lip. Since both the eCN and the precursors of another excitatory cell type, the cerebellar granule cells, express *Atoh1* but at different time points (E9-12 and after E13, respectively (Machold and Fishell, 2005)), the mice were provided with Doxycycline (Dox) starting at embryonic day 13.5 (E13.5) to shut down tTA activity after the eCN had extinguished *Atoh1* expression, thus restricting DTA expression to the eCN and sparing the granule cell precursors (Fig 2B). As shown in Fig. 2C-F, whereas control cerebella showed almost no cell death, in the cerebellum of the *Atoh1-tTA/+; Atoh1-Cre*/+; *Tigre*^*Dragon-DTA/+*^ mice given Dox at E13.5 onwards (*AC-eCN-DTA* model, AC designates *Atoh1-Cre* and eCN designates eCN-expression using *Atoh1-tTA*) ectopic cell death (marked by TUNEL staining) was detected specifically in the Nuclear Transitory Zone (NTZ), the region where the precursors of the eCN first migrate to after leaving the rhombic lip. tdTom^+^ cells were detected in the cells expected to express *Atoh1-tTA* including the NTZ, granule cell precursors in the external granule cell layer (EGL) and in *Atoh1*-derived precerebellar hindbrain nuclei (Fig. 2E). Quantification of cell death revealed an 8.85±3.35 fold increase in TUNEL^+^ particles in the NTZ in *AC-eCN-DTA* animals compared to their control littermates (*Tigre*^*Dragon-DTA/+*^) at E13.5 (n=3 per condition, Fig. 2G). At P1, the number of eCN, revealed by MEIS2 nuclear immunostaining, was reduced by more than 50% in *AC*-*eCN-DTA* cerebella as compared to controls (Fig. 2H-J).

**Figure 2.**
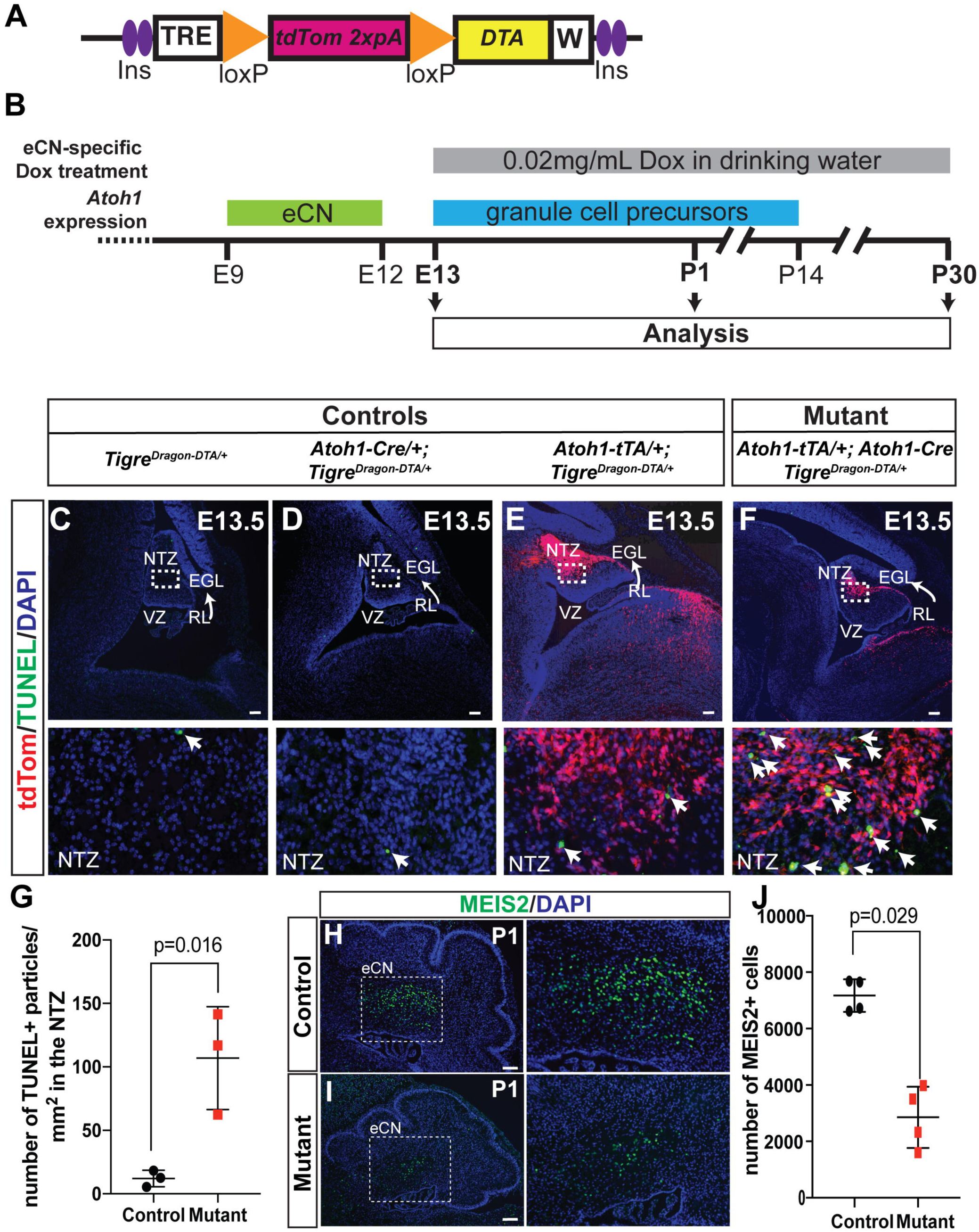
A *Tigre*^*Dragon-DTA*^ model to generate acute cell/tissue-specific cell ablation. A, B) Schematic showing *Tigre*^*Dragon-DTA*^ allele and experimental plan. C-F) Sagittal sections through the cerebellum of three types of E13.5 control embryos (*Tigre*^*Dragon-DTA*/+^, *Atoh1-Cre/+; Tigre*^*Dragon-DTA*/+^ and *Atoh1-tTA/+; Tigre*^*Dragon-DTA*/+^) and an *AC*-*eCN-DTA* embryo in the absence of Dox. The boxed areas are shown magnified in the lower panels. NTZ, nuclear transitory zone; VZ, ventricular zone; RL, rhombic lip; EGL, external granule layer. Long arrows show the direction of the tangential migration as the cells exit the RL. Arrowheads in the magnified boxes point to apoptotic cells (TUNEL^+^). (G). Quantification of cell death in the NTZ identified by MEIS2 staining on adjacent sections (not shown) of E13.5 embryos as measured by the number of TUNEL^+^ particles. *Tigre*^*Dragon-DTA*/+^ animals were used as controls (n=3/genotype, p=0.016, Mann-Whitney unpaired test). (H, I) Sagittal sections through the cerebellum at P1, showing reduced numbers of MEIS2^+^ eCN in *AC*-*eCN-DTA* mutants compared to a *Tigre*^*Dragon-DTA*/+^ control. (J) Quantification of MEIS2^+^ eCN number per half P1 cerebellum (every other section). *Tigre*^*Dragon-DTA*^/+ animals were used as controls (n=4/genotype, p=0.029, Mann-Whitney unpaired test). Scale bars = 100 µm.

One complication of using *Atoh1*-driven transgenes to target cells in the cerebellum is that they are also expressed in many precerebellar neurons in nuclei of the hindbrain at E10-13, as well as ectopic expression elsewhere in the brain and expression in other organs. Therefore, we used an intersectional approach to limit the expression of DTA to cells in the cerebellum and only a few nuclei in the isthmus in the midbrain-hindbrain junction (Machold and Fishell, 2005) by incorporating our *En1*^*Cre*^ knock-in line (Kimmel et al., 2000). This line expresses Cre in the midbrain and rhombomere 1 of the hindbrain starting at E8 and thus includes the rhombic lip (Fig. 3A). By using *Atoh1-tTA/+; En1-Cre; Tigre*^*Dragon-DTA/+*^ mice and providing Dox starting at E13.5 to shut down tTA activity in granule cell precursors (see Fig. 2B), we restricted DTA expression primarily to the eCN (Fig. 3B-E, *EC*-*eCN-DTA* model, EC designates *En1*^*Cre*^). Analysis of E13.5 embryos before Dox was administered showed ectopic cell death specifically in the region of the rhombic lip where cells first turn on the *Atoh1-tTA* transgene in *EC*-*eCN-DTA* mice (marked by TUNEL staining, Fig. 3B-E). Since *En1*^*Cre*^ turns on earlier than *Atoh1-Cre* in the eCN precursors, cell death was observed in the rhombic lip rather than in the NTZ (where it was seen in *AC-eCN-DTA* embryos). Quantification of cell death revealed a 14.20 ± 2.43 fold increase in TUNEL^+^ particles in the rhombic lip region of the *EC*-*eCN-DTA* embryos compared to controls (*Tigre*^*Dragon-DTA*/+^ or *Atoh1-tTA/+; Tigre*^*Dragon-DTA*/+^, n=3/condition, Fig. 3F). Furthermore, unlike in *AC*-*eCN-DTA* embryos, no tdTom^+^ cells were detected in the NTZ, consistent with the majority of eCN being ablated before tdTom is induced since Cre is present before the *Atoh1-tTA* transgene is expressed (Fig. 3E). In addition, at E13.5 we observed tdTom^+^ cells in the hindbrain posterior to rhombomere 1 that belong to the Atoh1-lineage and do not express *En1* or Cre (Machold and Fishell, 2005) (Fig. 3D,E). At P30, the number of eCN, revealed by counting of the large NeuN^+^ neurons in the cerebellar nuclei was reduced by ∼60% in *EC*-*eCN-DTA* cerebella as compared to the controls (Fig. 3G). As a consequence of the embryonic killing of the eCN, the size of the cerebellum was reduced by approximately one half (Fig. 3H-J). The decrease in cerebellum size at P30, and thus growth during development, was greater with the intersectional *EC*-*eCN-DTA* model than with the *AC*-*eCN-DTA* model (Willett et al. 2019), likely because in the latter some eCN do not express DTA because *Atoh1-Cre* is only transiently expressed in eCN precursors and once it is expressed it must first delete the floxed *tdTom* gene in the *Tigre*^*Dragon-DTA*^ allele to allow DTA to be expressed and kill cells. Consistent with this, in *AC*-*eCN-DTA* E13.5 embryos tdTom^+^ cells were detected in the NTZ and granule cell precursors (Fig. 2). Comparison of the intersectional approach (*Atoh1-tTA* and *En1*^*Cre*^) with the double *Atoh1*-driven transgenes (*Atoh1-tTA* and *Atoh1-Cre*) demonstrates the unique utility of each approach, with the *En1* and *Atoh1* intersectional approach achieving more precise cell targeting of DTA, and also earlier and therefore more widespread killing of eCN. Furthermore, analysis of *Tigre*^*Dragon-DTA/+*^ and *Atoh1-tTA/+; Tigre*^*Dragon-DTA/+*^ mice demonstrates that the transgene does not have leaky expression of DTA when the *tdTom* STOP sequence is present, while analysis of either of the control transgenic embryos with *Cre* alone (*Atoh1-Cre* or *En1*^*Cre*^) demonstrates lack of leaky expression of the recombined allele in the absence of tTA (Fig 2. C-E and Fig. 3 B-D).

**Figure 3.**
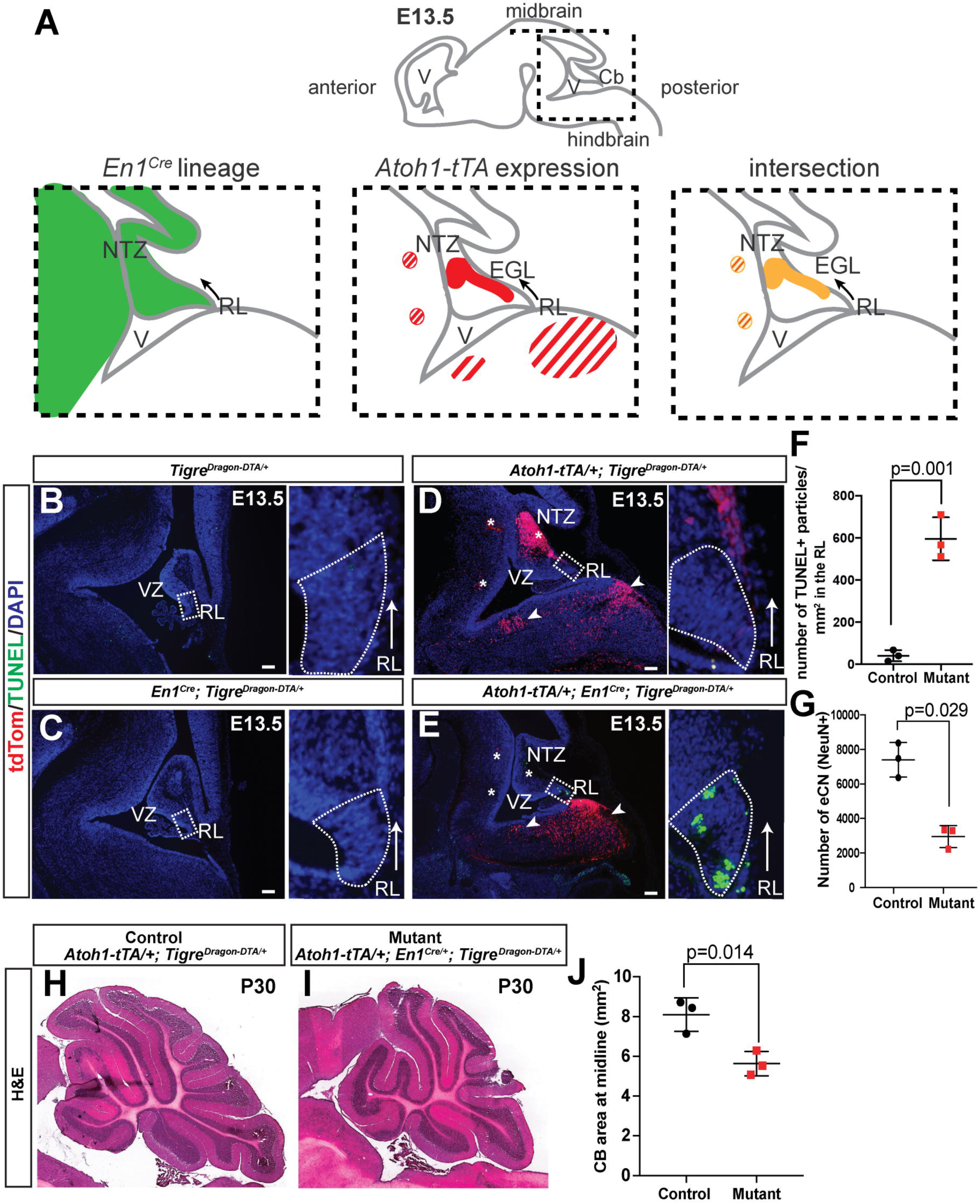
Intersectional capabilities of the *Tigre*^*Dragon-DTA*^ model. A) Schematic of *En1-*lineage (based on (Sgaier et al., 2007)) and *Atoh1* expression (based on (Machold and Fishell, 2005)). Striped regions represent hindbrain nuclei that are outside of the cerebellum. B-E) Representative sagittal sections showing the cerebellar anlagen at E13.5 in control embryos (either *Tigre*^*Dragon-DTA/+*^, *En1*^*Cre/+*^; *Tigre*^*Dragon-DTA*/+^, or *Atoh1-tTA/+; Tigre*^*Dragon-DTA*/+^) and *EC*-*eCN-DTA* embryos, in the absence of Dox. Asterisk points to the loss of tdTom^+^ cells in the NTZ and the two hindbrain nuclei targeted by the intersectional approach and the arrowhead points to the hindbrain nuclei that are not affected. The boxed areas are shown magnified in the right panels, rotated counter-clockwise. Insets show cell death (TUNEL^+^) in the rhombic lip region (RL). Arrows show the direction of the tangential migration as the cells exit the RL. The boxed areas are shown magnified in the right panels and were rotated counter clockwise. F) Quantification of cell death in the RL of E13.5 embryos detected by TUNEL (n=3/genotype, p=0.001, Mann-Whitney unpaired test). G) Quantification of large NeuN^+^ cells from every other sagittal section from half P30 cerebella shows the number of excitatory cerebellar nuclei in *EC-eCN-DTA* animals as compared to controls (n=3 for both genotypes, p=0.029, Mann-Whitney unpaired test). H-I) Representative images of the P30 cerebella of *EC*-*eCN-DTA* (I) and littermate control (H). J) Quantification of cerebellar area of midline sections shows reduction in cerebellar area after eCN ablation. Scale bars = 100 µm. *Tigre*^*Dragon-DTA*/+^ or *Atoh1-tTA/+; Tigre*^*Dragon-DTA*/+^ animals were used as controls for the quantifications shown in F, G and J. NTZ, nuclear transitory zone; VZ, ventricular zone; V, ventricle; RL, rhombic lip; EGL, external granule layer.

To further demonstrate the utility of the *Tigre*^*Dragon-DTA*^ line we bred it with mice bearing a left-lateral plate mesoderm specific *Pitx2-Cr*e (Shiratori et al., 2006) and either one of two possible cartilage-specific (r)tTAs: *Col2a1-tTA* (Rosello-Diez et al., 2018) (for E15.5 experiment, no Dox added) or *Col2a1-rtTA* (Posey et al., 2009) (for P1 experiment, Dox added at E12.5) to achieve unilateral ablation in the cartilage of the left limbs (*PC-Cart-DTA* model, PC designates *Pitx2-Cr*e and Cart designates cartilage-specific expression using *Col2a1-(r)tTA;* Fig. 4A). At E15.5 (i.e. ∼3 days after activation), TUNEL staining revealed increased cell death in the left cartilage compared to the right one (Fig. 4B), which was also seen at P1 (Fig. 4D). Quite strikingly, maintenance of the injury until P0 led to the generation of acellular gaps in the left cartilage (dashed lines, Fig. 4C, n=3). This injury led to a significant left-right limb asymmetry at P0, with the left bones 10-15% shorter than the contralateral ones (Fig. 4E).

**Figure 4.**
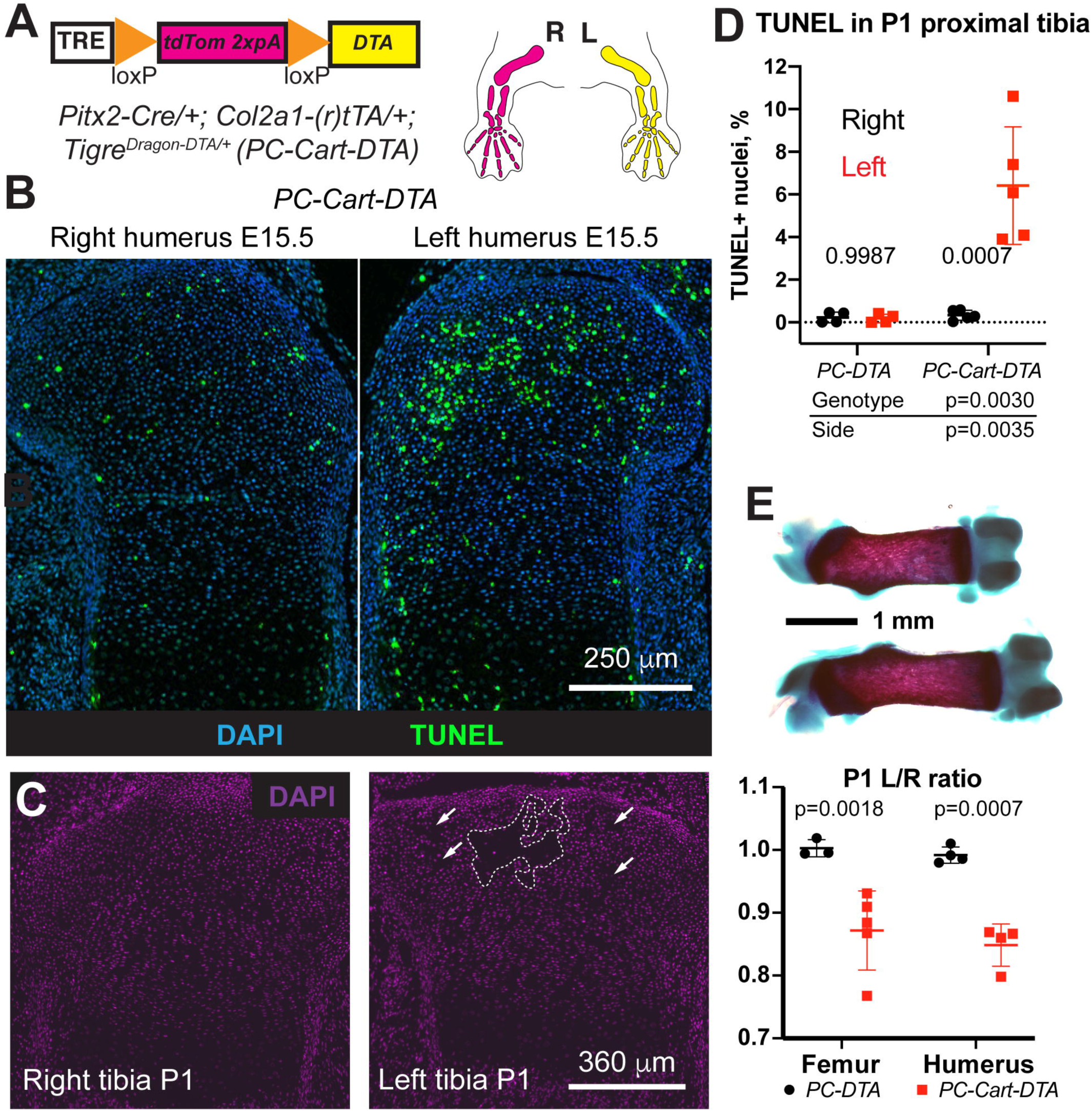
*Tigre*^*Dragon-DTA*^ allows generation of structural damage and reduced local cell density due to cell death in tissues with low cell motility such as cartilage. A) Schematic of the unilateral cartilage-specific cell ablation model. *Col2a1-tTA* (Tet-Off) starts to be expressed in the limbs atm ∼E12.5, the same stage at which Dox was provided in the Tet-On model to activate *Col2a1-*driven rtTA (and maintained until collection). B) TUNEL staining on E15.5 proximal humerus sections shows increased apoptosis in the left cartilage as compared to the right. C) One week of continuous DTA expression leads to the generation of acellular gaps in the left cartilage (dashed lines and arrows). D) Quantification of TUNEL^+^ cells in cartilage at P1 (n=4 *PC-DTA* and 5 *PC-Cart-DTA*). 2-way ANOVA results are shown. The bottom table shows the p-values for the different variables. p-values within the graph are for Sidak’s multiple comparisons test. E) Cell death in the cartilage leads to impaired left-bone growth as compared to the contralateral one at P1 (representative femora shown in top panel). Bone length ratios were analyzed by a mixed effects model (pGenotype <0.0001), as some samples lacked matching femur-humerus measurements. p-values for Sidak’s multiple comparisons post-hoc test are shown on top.

With toxins as powerful as DTA, it is important to target only the population of interest, which is one of the main advantages of our intersectional system. Indeed, *Pitx2-Cre* is expressed not only in the left lateral plate mesenchyme, but also in some regions of the heart (Shiratori et al., 2006), which can lead to unwanted effects. For example, if expression of diphtheria toxin receptor is activated by *Pitx2-Cre*, injection of diphtheria toxin leads to cell death not only in the left limb mesenchyme, but also in the heart, affecting survival of the experimental animals (Supplementary Fig. 1A and (Rosello-Diez et al., 2017)). Importantly, with our intersectional system almost no cell death was detected in the hearts of *PC-DTA* controls or Dox-treated *PC-Cart-DTA* embryos (Supplementary Fig. 1B-D), confirming the validity of the intersectional approach and lack of DTA expression in the absence of active (r)tTA. In summary, combining our *Tigre*^*Dragon-DTA*^ line with different pairs of the available *Cre* and *(r)tTA* lines provides a valuable resource to selectively ablate cells with high spatiotemporal precision.

### Unilateral misexpression of Connective Tissue Growth Factor in the growing skeletal elements leads to bone asymmetry

We next searched for a GOI that could provide a versatile tool for transiently manipulating a variety of biological processes, such as development, homeostasis and repair. We reasoned that a molecule capable of interacting with several of the most common signaling pathways would be an ideal candidate. Connective tissue growth factor (CTGF) has been described to exert such a pleiotropic effect (Takigawa, 2013). Indeed, CTGF can interact with several signaling receptors, is considered as a hub in a network of molecular interactions, and plays a role in the harmonization of cartilage and bone development (Kubota and Takigawa, 2015), cardiovascular function (Ponticos, 2013), fibrosis, tendon healing (Tarafder et al., 2017), etc. We thus generated a *Tigre*^*Dragon-Ctgf*^ mouse model (see Methods) and combined it with mice bearing left-lateral plate mesoderm specific *Pitx2-Cre* (Shiratori et al., 2006) and a cartilage-specific *Col2a1-tTA* (Tet-Off) (Rosello-Diez et al., 2018) to achieve unilateral expression of CTGF in the cartilage of the left limbs (*PC-Cart-Ctgf* model, Fig. 5A). By P0-P1, whereas the expression of *Ctgf* in the distal cartilage of the right *PC-Cart-Ctgf* radius was very similar to the expression in WT animals (Fig. 5B-C), the expression in the left *PC-Cart-Ctgf* element was more widespread (Fig. 5C). Interestingly, the ectopic expression of CTGF led to reduced growth of the left bones as early as embryonic day (E) 17.5, including deformation of the radius and ulna at or after postnatal day (P) 5 (Fig. 5D). While mechanistic studies will be reported elsewhere, we are currently investigating whether this phenotype is related to increased vascular invasion and altered osteoclast activity, similar to that reported upon increased VEGF activity (Maes et al., 2010). Given its strong effects in a plethora of tissues and processes, we expect the *Tigre*^*Dragon-Ctgf*^ mouse line to become a valuable tool for life scientists working on diverse topics.

**Figure 5.**
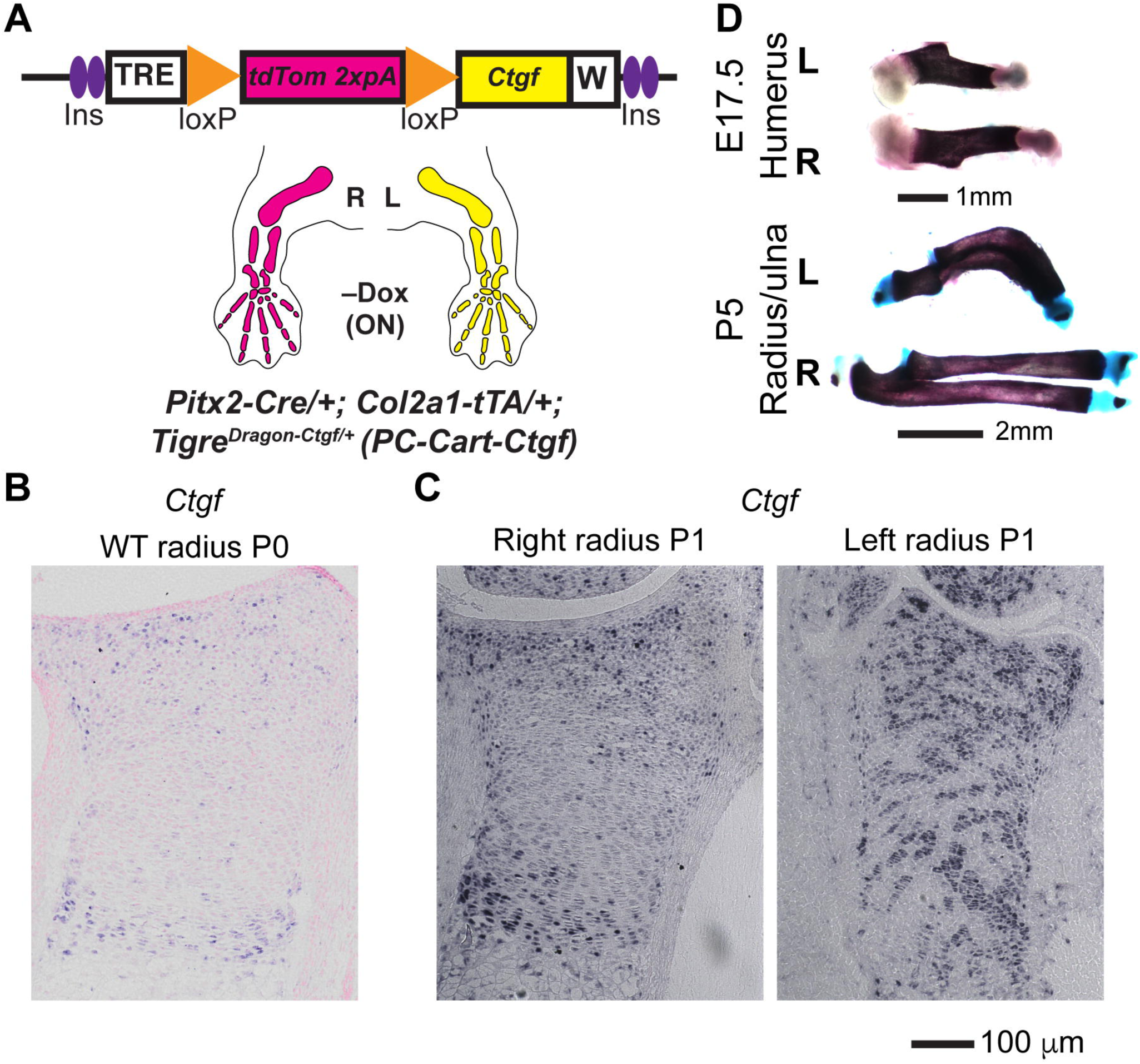
Unilateral overexpression of connective tissue growth factor leads to bone asymmetry. A) *PC-Cart-Ctgf* model to achieve misexpression of *Ctgf* in the cartilage of the left limb. Note that *Col2a1-tTA* (Tet-Off system) is used in this model, such that tTA is active without Dox (starting at ∼E12.5). B, C) *In situ* hybridization for *Ctgf* mRNA in the distal radius cartilage from WT (B) and *PC-Cart-Ctgf* mice (C) at P0-P1 (n=3). D) Representative examples of skeletal preparations of left and right *PC-Cart-Ctgf* bones at the indicated stages (n=7 at E15.5, 6 at E17.5, 4 at P1, 5 at P5-P7).

### A model for reversible connective tissue growth factor misexpression in the growing skeletal elements reveals different outcomes of transient vs. permanent expression

Besides its intersectional quality, a key aspect of the *Dragon* system is its reversibility. As a second proof-of-principle for testing this application, we repeated the *PC-Cart-Ctgf* experiment and supplied Dox to shut down tTA expression at E17.5 (Fig. 6A), soon after the phenotype is obvious (Fig. 5D). As expected, *Ctgf* ceased to be overexpressed in the cartilage by P0-P1 (Fig. 6B). Moreover, when the extent of asymmetry was compared between continuous and transient overexpression, we found that the latter led to a much milder asymmetry than the former (Fig. 6C). The transient expression experiment thus suggests that there is a catch-up mechanism that allows limb growth to recover, and that would not have been discovered by irreversible misexpression, highlighting the utility of the new mouse lines.

**Figure 6.**
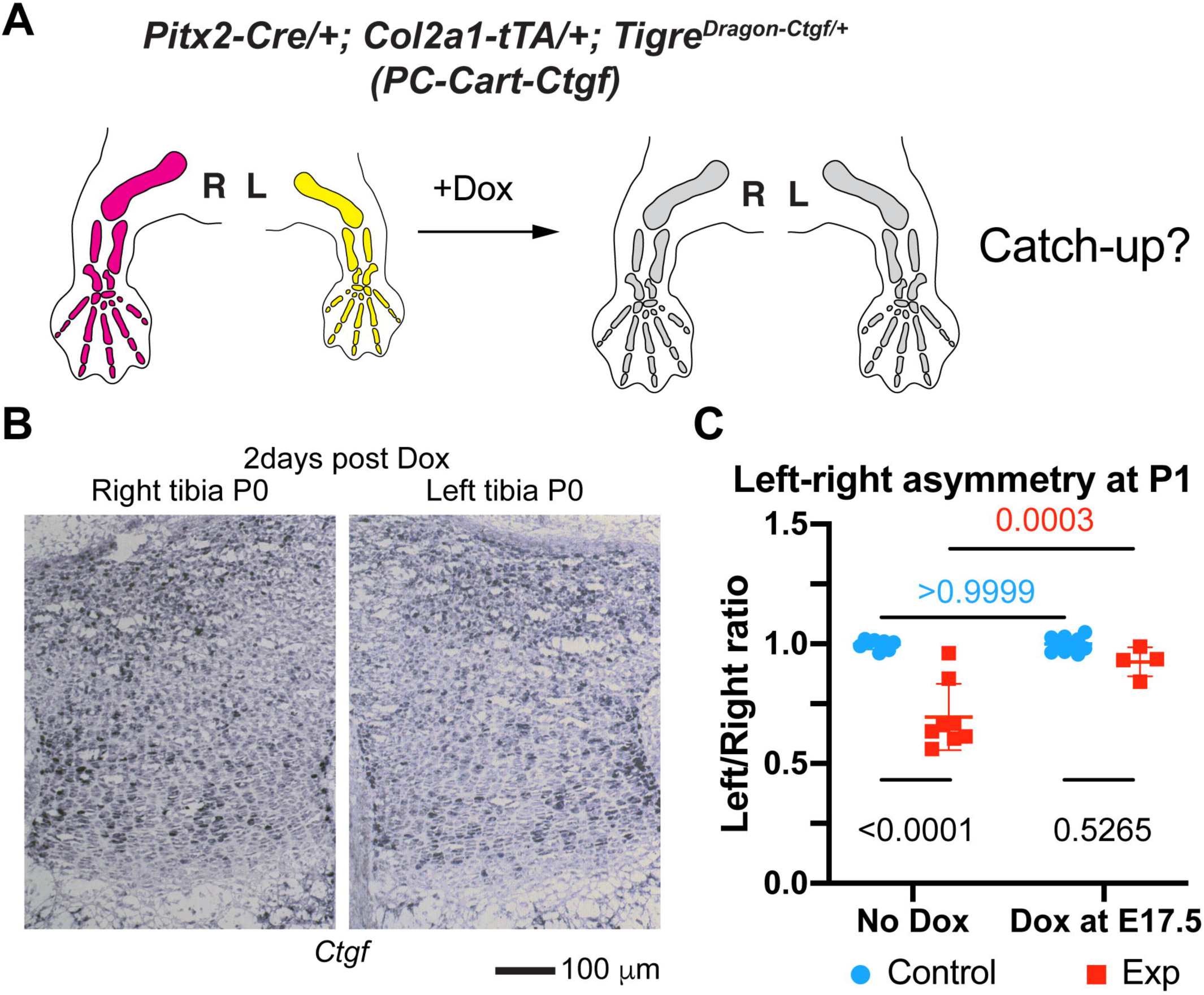
Different outcomes of transient vs. continuous expression of *Ctgf* in the cartilage of the growing bones. A) Schematics of the transient injury experiment. B) RNA in situ analysis of expression of *Ctgf* in the proximal cartilage of left and right tibia of *PC-Cart-Ctgf* P0 pups, 2 days after Dox administration at E17.5 (n=3). C) When transgenic *Ctgf* expression was shut down by Dox treatment in the *PC-Cart-Ctgf* model (Dox at E17.5, analysis at P0-P1), the asymmetry was not as extreme as in the case of continuous *Ctgf* misexpression (no-Dox condition). In the graphs, quantification of left/right ratio of bone length is shown for control (tTA^−^) and experimental animals (tTA^+^), treated or not with Dox. Analysis by 2-way ANOVA for Genotype and Treatment, p-values for Sidak’s multiple comparisons post-hoc test are shown. Since we used an internal ratio (left/right length), data from femora and humeri were pooled together. They were treated as independent samples because *Pitx2-Cre* has been shown to recombine with different efficiencies in forelimb and hindlimb (Rosello-Diez et al., 2018).

### The *Tigre*^*Dragon*^ lines are effectively activated by (r)tTA and Cre, and not leaky

We then performed in-depth characterization of the described lines, to ensure they were tightly and efficiently regulated in a variety of scenarios. In previous studies we showed that *TRE*-driven transgenes are highly expressed from the *Tigre* locus when insulator elements and a Woodchuck virus RNA stabilizing sequence are included (Madisen et al., 2015; Rosello-Diez et al., 2018). However, it has not been tested whether the TRE in the *Tigre*^*Dragon*^ lines drive expression of GOIs at a high level in the presence of other *TREs* that could compete for the transcriptional machinery and rtTA. This is important because one possible non-intersectional application of our system is to combine it with an (r)tTA and a TRE-driven Cre to achieve expression of both Cre and the GOI in a tissue during the time (r)tTA is activated. To address potential competition between TRE-driven transgenes, we generated mice bearing a *TRE-Cre* (Perl et al., 2002), our *Col2a1-tTA* allele–which initiates expression in the cartilage at around E12.5 (Rosello-Diez et al., 2018)–and the *Tigre*^*Dragon-p21*^ allele (*TC-Cart-p21* model, Fig. 7A,B). We analyzed the mice at postnatal day (P) 0 in the absence of Dox, such that the tTA was active for at least a week before analysis. In *Tigre*^*Dragon-p21*^ mice that lacked both *tTA* and *TRE-Cre*, no tdTom or p21 expression was detected in the target tissue (or other tissues), confirming that there is no spontaneous expression of the *Dragon* allele (Fig. 7C and C’ left). In the presence of only *tTA*, ectopic expression of almost only tdTom was found in the cartilage (Fig. 7C middle), and the intensity of the few p21^+^ cells was barely above background (Fig. 7C’ middle, D), revealing there is negligible read-through activity past the STOP cassette. Finally, when both *TRE-Cre* and *tTA* were present along with the *Tigre*^*Dragon-p21*^ allele, we observed only p21 expression, demonstrating a full switch from *Tigre*^*TRE*^-driven tdTom to p21 expression (Fig. 7C, C’, D). Since no cells expressed only tdTom in this genetic combination, this result indicates that the two *TRE-*driven alleles are both effectively induced.

**Figure 7.**
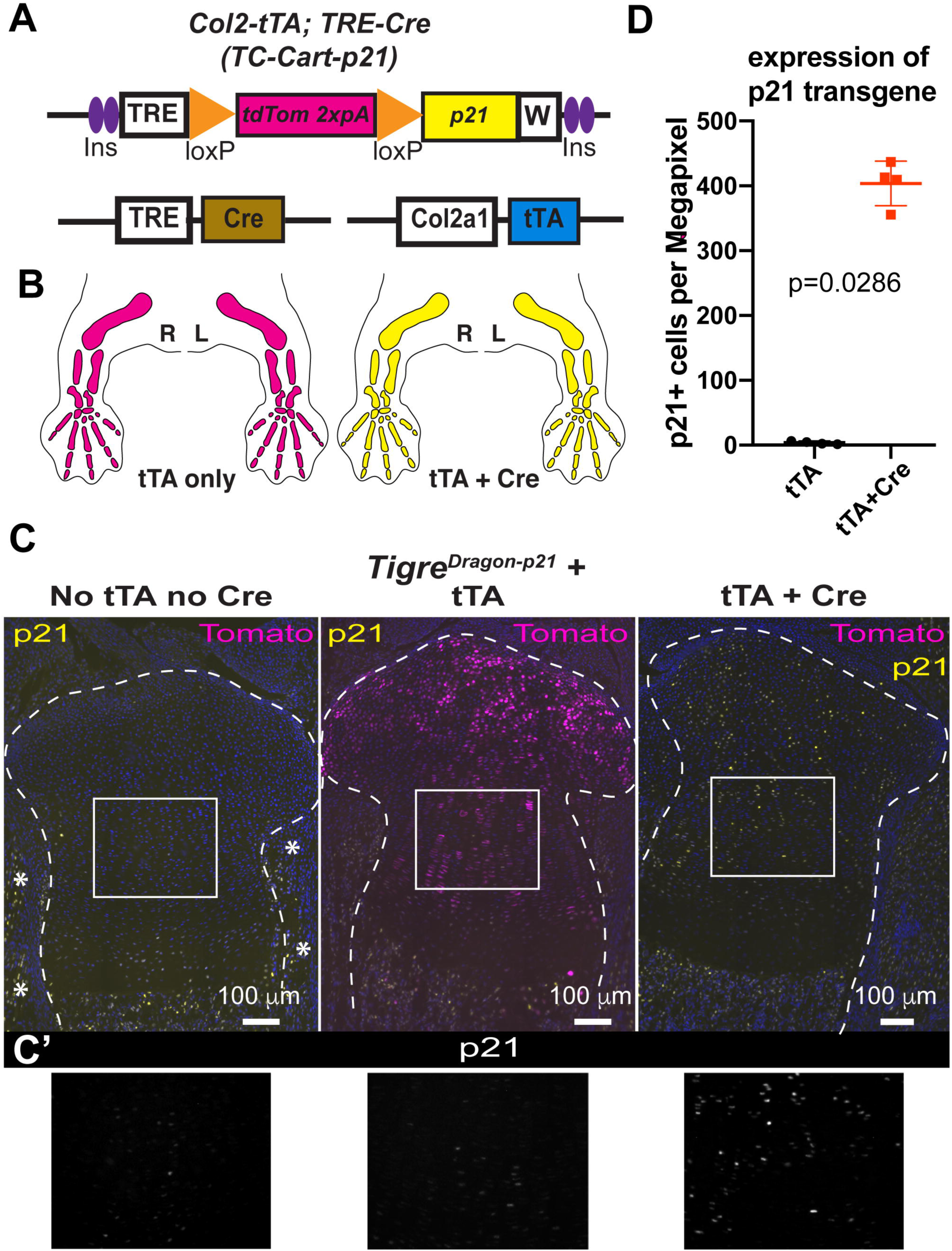
The *Dragon* lines are effectively activated by tTA and Cre, and not leaky. A) Depiction of the alleles that compose a *TC-Cart-p21* model (see text for details). B) Prediction of expression domains [color code as in (A)] in the presence of *Col2a1-tTA* but absence of the Cre allele (left), and when both Cre and tTA alleles are present along with *Dragon-p21* (right). R, L: right and left limbs, respectively. C) Representative micrographs of immunohistochemistry for the indicated proteins on proximal tibia sections from newborn littermates of the indicated genotypes (*Tigre*^*Dragon-p21*^ is common to all three). Dashed lines delimit the cartilage. Asterisks indicate endogenous expression of p21 in muscle and perichondrium. No Dox was provided for these experiments. C’) 1.8X magnification of boxed areas in C). Only p21 staining is shown. D) Quantification of p21 expression in control and experimental mouse sections (n=4 each). p-value for unpaired Mann-Whitney test is shown.

### A germline-recombined version of the *Tigre*^*Dragon*^ allele is controlled by (r)tTA only

In cases where intersectionality is not a priority, to avoid the extra breeding steps required to introduce a third allele such as *TRE-Cre*, we have also generated a germline-recombined version of the *Tigre*^*Dragon-p21*^ allele (*Tigre*^*TRE-p21*^ or *p21rec*, Fig. 8A) that only requires (r)tTA activity to express *p21*. To characterize this allele, we generated mice containing *Col2a1-rtTA, p21rec* and *Pitx2-Cre* (the latter for control purposes, *Cart-p21* model). As expected, when we provided Dox to induce rtTA activity at E12.5, since p21 expression does not require intersection with Cre activity, there was no significant left-right difference in the number of chondrocytes expressing p21 (Fig. 8B,C). Moreover, a higher percentage of chondrocytes expressed p21 in the limbs (∼80%, Fig. 8B,C) than in the left limb of the intersectional approach (50-60% in Dox-treated *Pitx2-Cre/+; Col2a1-rtTA; Tigre*^*Dragon-p21*^ mice, (Rosello-Diez et al., 2018)). This new *p21rec* strain and similar derivatives that could be generated from our *DTA* and *Ctgf Dragon* alleles will be useful to interfere with cell functions in a variety of contexts, either by using existing tissue-specific (r)tTA mouse lines or Cre-dependent (r)tTAs activated by tissue-specific Cre.

**Figure 8.**
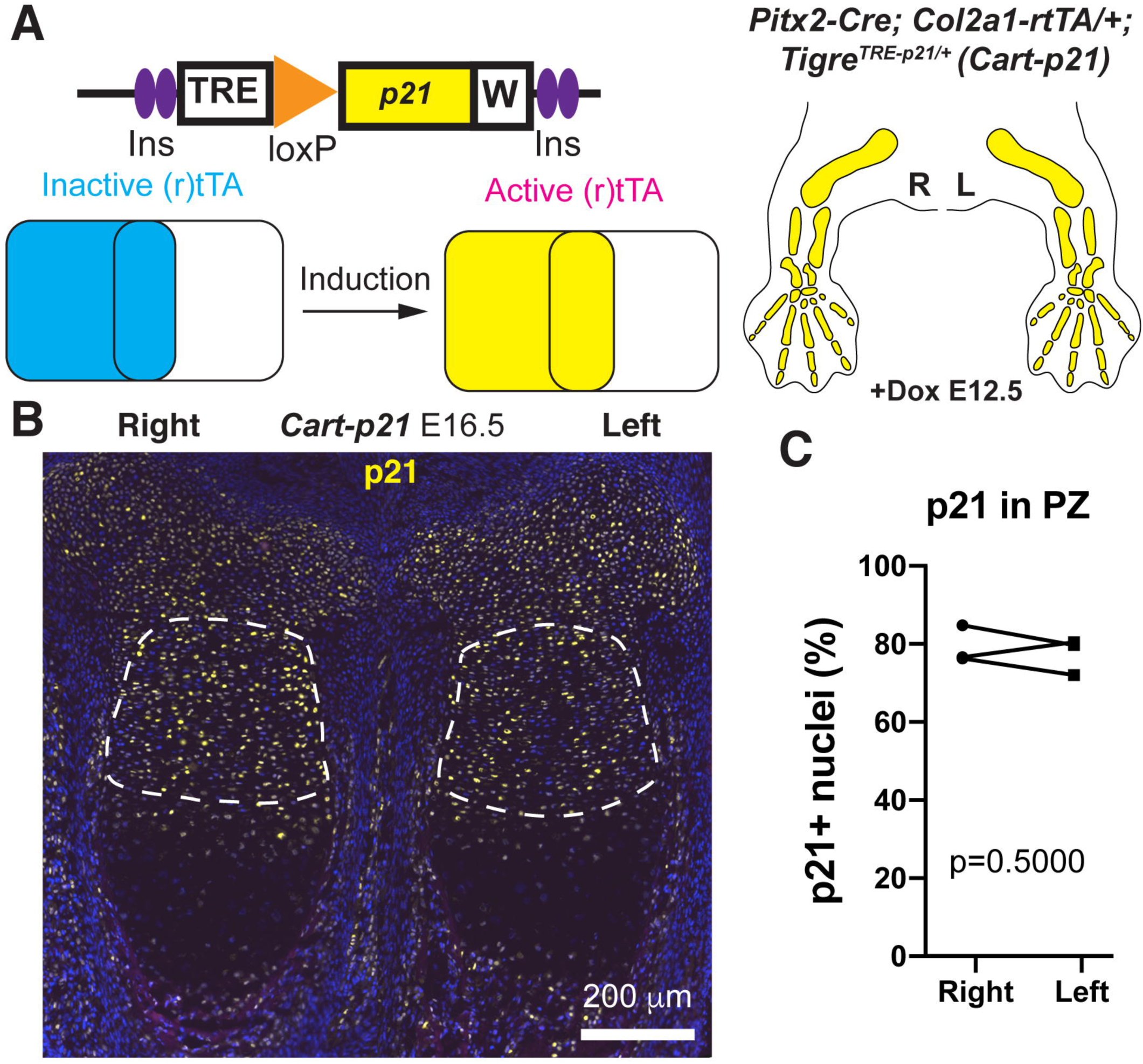
A germline-recombined version of the *Tigre*^*Dragon-p21*^ allele becomes responsive to active (r)tTA only. A) Depiction of the recombined allele (*p21rec*) and how it works in the presence of *Col2-rtTA* (*Cart-p21* model). Note that the crosses include *Pitx2-Cre* for control purposes, even though it is not necessary for p21 expression. B) Representative micrograph showing p21 expression in left and right proximal tibial cartilage at E16.5, 4 days after Dox administration. C) Quantification and statistical analysis of p21^+^ cells in the proliferative zone (PZ, delimited by dashed lines). p-value for Wilcoxon matched-pairs signed rank test is shown.

### The *Dragon* targeting vectors can be used for *in ovo* electroporation

Lastly, we tested the possibility of using the *Tigre*^*Dragon*^ tools in animal models other than the mouse. The chicken embryo has been a classical model for cell and developmental biology, immunology, virology and cancer that is experiencing a revival since the introduction of modern genetic and imaging tools (Kain et al., 2014; Rashidi and Sottile, 2009; Stern, 2005). *In ovo* electroporation enables the introduction of exogenous genetic constructs in multiple locations of the embryo, in order to perform gain- and loss-of-function studies, lineage tracing, live imaging, etc. (Morin et al., 2017; Scaal et al., 2004). Therefore, we electroporated the right half of the neural tube of chicken embryos at Hamilton-Hamburger (HH) stage 14-17 (Hamburger and Hamilton, 1992) with a mix of *pCAGGS-H2B-TagBFP* (to visualize electroporated cells), *pCAGGS-rtTA-IRES-Cre* (to constitutively express rtTA and Cre) and a *Tigre* targeting vector (*Ai62*) bearing a *TRE-loxSTOPlox-tdTomato* cassette (Madisen et al., 2015). 16h after electroporation, Dox was provided (1 µg per egg). 24 h after electroporation, we observed robust and consistent electroporation (blue signal, Fig. 9A) in the right neural tube of most surviving embryos, as well as strong red signal in the electroporated area of Dox-treated embryos (Fig. 9A, C, n=11). Embryos not treated with Dox did not show any tdTom signal (Fig. 9B, n=3).

**Figure 9.**
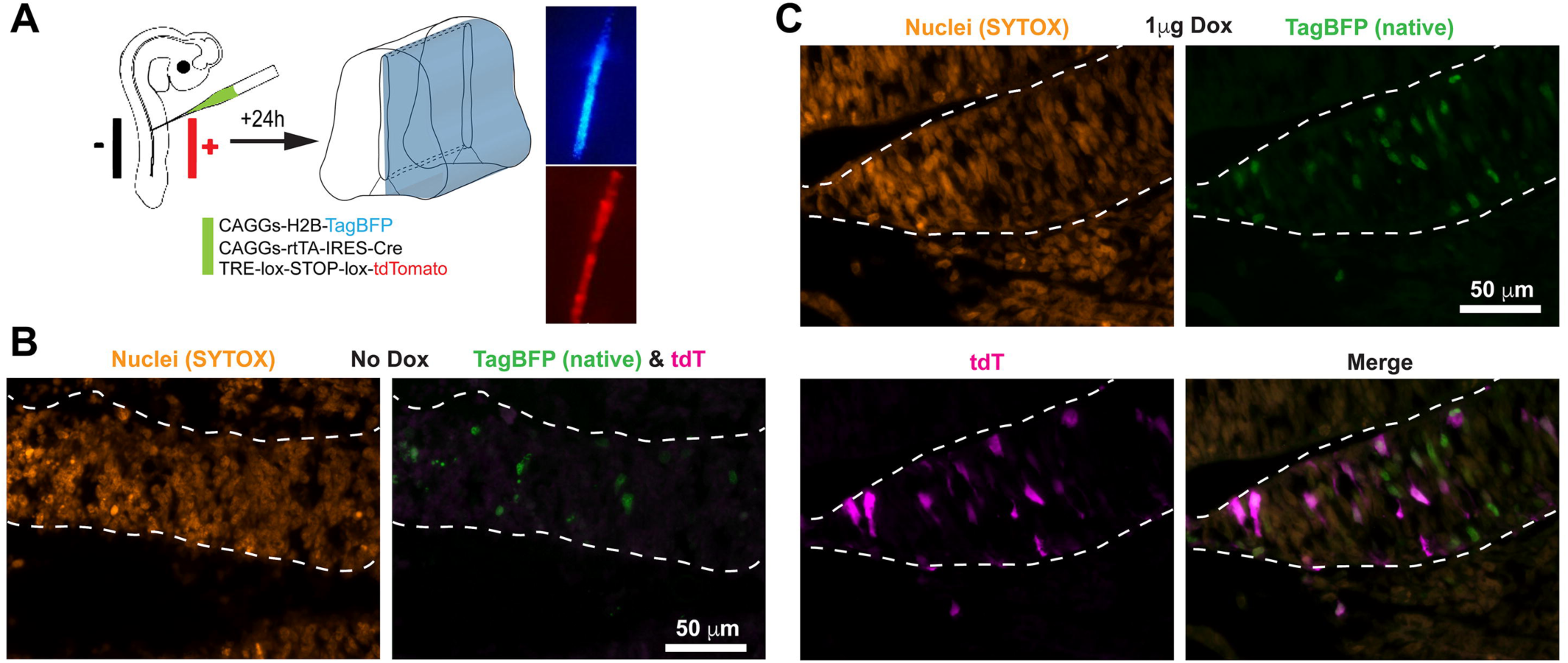
The *Dragon* vectors can be used for *in ovo* electroporation. A) Schematics of the electroporation procedure in the neural tube of chicken embryos (cartoon), and representative examples of the outcome (micrographs taken on a fluorescence dissecting microscope). B-C) Individual and merged channels of transverse neural tube sections (dashed lines) from electroporated chicken embryos, either untreated (B, n=3), or Dox-treated (C, n=11), 24 h after electroporation. In order to be able to visualize the native fluorescence of BFP, we used a green nuclear stain (SYTOX Green).

Lastly, we took advantage of chicken embryo accessibility to try an orthogonal approach. In this case, only pCAGGS-rtTA and the *Tigre* construct were electroporated, while the Cre was added as a fusion protein (TAT-Cre). The TAT peptide allows the Cre to cross membranes (Gitton et al., 2009), such that when beads loaded with TAT-Cre are implanted near the electroporated area, the Cre diffuses a few cell diameters away. The Cre then activates the GOI only in the electroporated cells that are close to the bead, and only in response to Dox (Fig. 10A,B, n=6). Control embryos electroporated with the full mix, but without bead (n=5, not shown), or implanted with PBS-soaked beads (Fig. 10C, n=5), or electroporated with a mix devoid of rtTA and implanted with TAT-Cre beads (Fig. 10D, n=5) did not show any tdTom^+^ cells.

**Figure 10.**
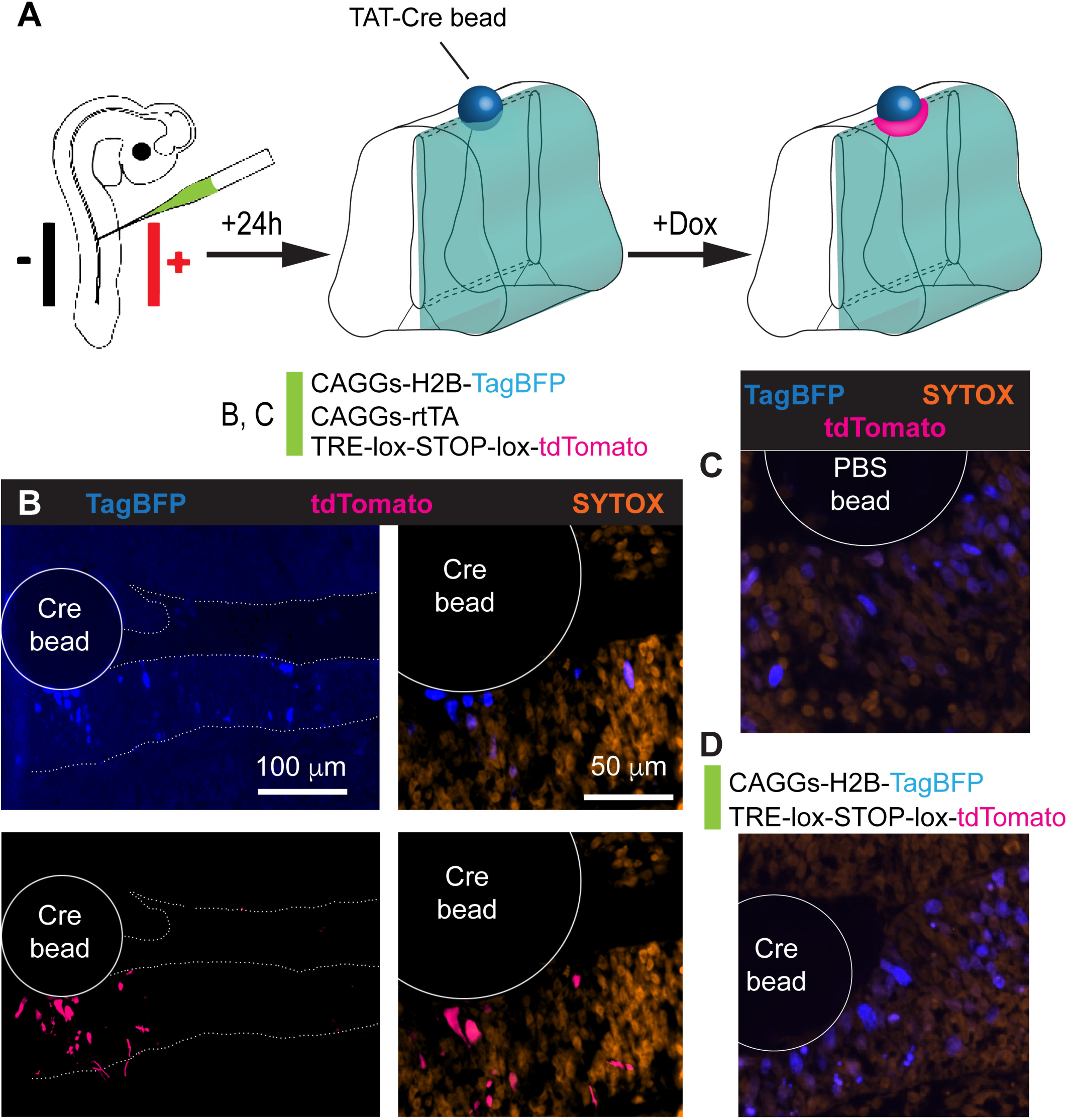
Orthogonal *Dragon-*driven gene misexpression combining electroporation and implantation of Cre-soaked beads. A) Schematics of the procedure of electroporation followed by bead implantation. B) Implantation of TAT-Cre beads (3 mg/ml) activates tdT expression in some of the electroporated cells (TagBFP^+^) close by, but not in cells farther than ∼100 microns from the bead. Native expression of TagBFP is shown. C, D) Control experiments with PBS-only beads and a full electroporation mix (C, n=5) and Cre-beads in the presence of an electroporation mix lacking the rtTA plasmid (D, n=5).

## DISCUSSION

In this report, we have expanded and further tested the repertoire of *Tigre*^*Dragon*^ intersectional tools and demonstrate their advantages for in vivo studies. Given that there are currently over 3,000 Cre / CreER lines (http://www.informatics.jax.org/downloads/reports/MGI_Recombinase_Full.html), hundreds of which are available in repositories, and >200 (r)tTA lines (almost half of them in repositories, see https://www.tetsystems.com/fileadmin/downloads/transgenic_mouse_lines/18-06-07_table_1_ta.pdf), the potential combinations with the tools we describe here are extremely numerous, allowing for very nuanced studies. As these numbers keep increasing, we anticipate that the *Tigre*^*Dragon*^ system will be useful to many developmental biologists and other life scientists across different fields. The three main mouse lines described here (*Tigre*^*Dragon-p21*^, *Tigre*^*Dragon-DTA*^ and *Tigre*^*Dragon-Ctgf*^) are being imported by the JAX repository (stocks # 034777-034779) to make them available for the scientific community. It should be noted that these lines can be maintained in homozygosity with no apparent phenotype, although after several generations of inbreeding some of the homozygous animals show reduced fertility. Our labs mostly maintain these lines in an outbred SwissWebster background. No parent-of-origin effects have been observed for these alleles. The DNA constructs are in the process of being submitted to Addgene (#140894-140896).

### Tigre^Dragon-DTA^

We have shown, both in cerebella and limbs, that the *Tigre*^*Dragon-DTA*^ line can be used for specific ablation of particular cell populations, restricted to a temporal window of interest. Coupled to the vast number of possible combinations of *Cre* and *(r)tTA* lines, the *Tigre*^*Dragon-DTA*^ strain will be a valuable resource to ablate cells at any developmental stage and location, allowing for systemic and local responses to cell loss to be studied, including the activation of signaling pathways in response to altered cell density and regenerative mechanisms to replenish cells that are lost.

### Tigre ^Dragon-Ctgf^

CTGF is a pleiotropic molecule, which makes it difficult to elucidate its specific roles by traditional overexpression approaches in which several locations and cell types are affected. Conversely, the intersectionality of our approach enables targeting more precisely to a defined region, during a time window of interest. This approach will enable the elucidation of tissue-specific roles of CTGF, which should positively impact on regenerative medicine applications, tissue engineering and developmental biology research.

### Avian experiments

Our results indicate that the plasmids we have generated for mouse ES cell targeting can be directly used for efficient *in ovo* electroporation, opening the door for precise gene manipulation in other animal models. The use of tissue-specific rather than ubiquitous promoters would make the system more versatile in the future. The orthogonal approach we describe using TAT-Cre enables a number of experimental approaches with high spatial and temporal precision. For example, after electroporation of the *rtTA* and the *Dragon* construct at an early stage, one can implant Cre-loaded beads in a very localized region of the embryo when it is still accessible, and defer Dox administration to a later time point such that the GOI can be activated even if the embryo has sunk or turned in the egg.

### Comparison with existing systems

Our system complements some of the existing sophisticated mouse genetic tools developed by the community over the last few years. For example, Belteki et al. (Belteki et al., 2005) combined tissue specific Cre with Cre-inducible rtTA and a TRE-controlled transgene to achieve conditional, inducible and reversible transgene expression. That system however is not intersectional, since the specificity depends only on the promoter driving Cre. Using the same number of transgenes (three) our strategy achieves the additional feature of refined spatial control. A similar system reduced the number of transgenes by making the rtTA and TRE-controllable cassettes part of a bicistronic cassette (Haenebalcke et al., 2013). However, the cassette is targeted to the *Rosa26* locus, where the TRE is prone to epigenetic inactivation (Godecke et al., 2017). Depending on the specifics of the study being undertaken, our tools should allow approaches that were technically hindered before. With regards to more recent tools involving CRISPR/Cas9 approaches, overexpression of endogenous loci can be achieved via a nuclease-defective Cas9 fused to a transactivator domain in a structure-optimized manner (Konermann et al., 2015). This strategy could be used to achieve results similar to ours, if Cas9 were under the control of TRE with the sgRNAs in tandem, as recently described (Jo et al., 2019). However, such a strategy would not be intersectional, nor would it allow for misexpression of genes not present endogenously, such as DTA or genes harboring coding mutations related to human disease. In summary, our approach fills a new niche in the ecosystem of genetic tools, thus enabling new types of studies.

### Limitations and future directions

One potential limitation of our vectors is that whereas the constructs work very well for transfection *in vitro* using HEK cells (not shown) and for chicken embryo electroporation, they are probably not suitable for efficient viral transduction, as they tend to be large (>10 kb) due to the inclusions of insulators and RNA-stabilizing sequences.

Moreover, with the current system it is not possible to trace the lineage of the cells where the intersectional activation took place, as the transgene stops being expressed when the (r)tTA is shut down or the cells transition to another fate where (r)tTA expression is lost. Future versions could be designed to express Flp recombinase in addition to the GOI, and combined with Flp-reporters to indelibly label the lineage of the targeted population.

Lastly, we envision that the use of CreER instead Cre will introduce an additional level of control. For example, the CreER could be used to prime a stem cell population shortly after an injury, such that the descendants lose the loxSTOPlox cassette. In this scenario, only the descendants that differentiate into an (r)tTA-expressing cell population of interest would be targeted, as opposed to the whole set of descendants.

## MATERIALS AND METHODS

### Reagents and resources

See Supplementary Table 1.

### Generation and maintenance of mouse lines

The generation of the *Tigre*^*Dragon-p21*^ mouse line was as described (Rosello-Diez et al., 2018). To generate the *Tigre*^*TRE-p21*^ mouse line (aka *p21rec*), *Tigre*^*Dragon-p21*^ males were crossed with germline Cre-expressing females to delete the floxed cassette. The Cre allele was subsequently eliminated during the breeding to generate *Tigre*^*TRE-p21*^ homozygous animals.

To generate the *Tigre*^*Dragon-Ctgf*^ mouse line, the *NruI*-STOP-loxP-tdTomato-*SnaBI* fragment in the *Ai62(TITL-tdT) Flp-in* replacement vector (Madisen et al., 2015) was replaced by a custom *NruI*-*tdTomato-STOP-loxP*-*MluI-HpaI-SnaBI* cassette, to generate an empty DRAGON vector. A PCR-amplified Kozak-*Ctgf* cassette was subsequently cloned into the *MluI* and *SnaBI* sites to generate the *DRAGON-Ctgf* vector. This vector was then used for recombinase-mediated cassette exchange into *Igs7-*targeted G4 ES cells (Madisen et al., 2015). Two successfully targeted clones (#1 and #2) were injected into C2J blastocysts to generate chimeras, obtaining 11 chimeric males that survived (out of 30 animals born) with 25-85% chimaerism. One male from clone #1 and three males from clone #2 were crossed to Swiss Webster mice (Charles River) to assess germline transmission. Only clone #2 was successfully transmitted and used to establish the new mouse line.

A similar process was used to generate the *Tigre*^*Dragon-DTA*^ mouse line, with the only difference being that an attenuated version of DTA (tox176) was used to PCR-amplify the Kozak-DTA cassette. Two successfully targeted clones (#8 and #13) were injected into C2J blastocysts to generate chimeras, obtaining 34 chimeric males that survived (out of 52 animals born) with 35-100% chimaerism. Four males from clone #8 and two from clone #13 were used to assess germline transmission and establish the new mouse line.

All mouse lines were maintained on an outbred Swiss Webster background. *Atoh1-tTA, En1*^*Cre*^ and *TetO-Cre* were genotyped as described (Kimmel et al., 2000; Perl et al., 2002) (Willett et al., 2019). The DNA primers and programs used for genotyping the rest of alleles and transgenes are shown in Supplementary Table 2.

### Animal experiments

All mouse work was carried out according to institutional guidelines after obtaining their approval (Sloan Kettering Institute IACUC and Monash University Animal Ethics Committee). For time-mated crosses, vaginal plugs were checked in the mornings, and noon of the day the plug was detected was considered E0.5. When Dox was needed, it was added to the drinking water. To activate *Col2a1-rtTA*, the final concentration was 1 mg/ml, including 1% sucrose to increase palatability. To inactivate *Col2a1-tTA* and *Atoh1-tTA*, the final concentration of Dox was 0.02 mg/ml.

Upon reception, chicken eggs were kept at 18°C for up to 2 weeks. Two and half days prior to the experiment, they were transferred to a watered incubator at 37°C with forced ventilation.

### In ovo electroporation

*In ovo* electroporation was performed as described (Blank et al., 2007). Briefly, on the day of the experiment, eggs were laid out horizontally; 3 ml of albumin were removed from the blunt end of each egg, and a window was opened on top with blunt forceps. A few drops of Ringer’s solution containing Penicillin-Streptomycin (Gibco 15140122, diluted 1/50) were pipetted on top, and the membranes removed with a hooked needle. Indian ink (drawing ink A, Pelikan #201665, diluted 1/100 in Ringer’s solution) was injected under the vitelline membranes to provide contrast. A pulled glass capillary (Harvard apparatus GC120T-10) and a mouth pipette were used to inject into the caudal neural tube the electroporation mix (1 µg/µl each plasmid in injection solution: high viscosity carboxymethylcellulose 0.33% (Sigma); Fast Green 1% (Sigma); MgCl_2_ 1 mM in PBS). Custom-made electrodes (using 0.5-mm wide platinum wire for the positive electrode and a tungsten wire for the negative one) were placed flanking the embryo, and a pulse generator (Intracel TSS20 with current amplifier) used to deliver 3 square pulses (27 V, 30 ms ON, 100 ms OFF). After electroporation, 200 µl of Doxycycline solution (5 ng/µl in Ringer’s) were micropipetted underneath the embryo. Eggs were then sealed with tape and incubated for 24h.

### Bead implantation

Affigel blue beads were soaked at least 30 min in a solution of TAT-Cre (0.5 mg/ml in PBS), and rinsed in PBS before grafting. Implantation was performed as described (https://bio-protocol.org/e963), using a tungsten needle to open up a small section of the neural tube, and forceps Dumont #5 to insert the bead inside the socket.

### Cloning of pCAGGS-rtTA-IRES-Cre

The vector for constitutive expression of rtTA and Cre was built via Gibson cloning using Addgene #20910 (digested *SmaI-PacI*) as backbone, and the following fragments: an *IRES* sequence amplified from Addgene #102423 using primers CTTGACATGCTCCCCGGGTAACCCCTCTCCCTCCCCCCC and GCTCCATTCATTTATCATCGTGTTTTTCAAAGGAAAACCACG, and a Cre sequence amplified from genomic DNA of a *Cre*^*+*^ mouse with primers AAACACGATGATAAATGTCCAATTTACTGACCG and ATGCTCTCGTTAATCTAATCGCCATCTTCCAG. The molecular assembly was done with the NEBuilder HiFi DNA Assembly cloning kit following manufacturer instructions.

### Sample collection and processing

#### Mouse limbs

Upon embryo or pup collection, the limbs were dissected out in cold PBS, skinned and fixed in 4% PFA for 2 days. No decalcification was done for samples P1 or younger. After PBS washes, the limbs were cryoprotected in PBS with 15% sucrose and then equilibrated at 37°C in PBS with 15% sucrose and 7.5% porcine skin gelatin (Sigma) until they sank. The limbs were oriented inside the wells of multi-welled plates and cooled down until solid, to then flash-freeze them using a dry-ice cold bath of isopentane (Merck). The cryoblocks were sectioned at 7 µm on a cryostat (Leica, CM3050S) onto Superfrost slides.

#### Mouse brains

The brains of all embryonic stages and P1 pups were dissected in cold PBS and immersion fixed in 4% PFA for 24 hours at 4^°^C. P30 animals were transcardially perfused with PBS followed by 4% PFA, and brains were post-fixed in 4% PFA overnight. Specimens for cryosectioning were placed in 30% sucrose/PBS until they sank, embedded in OCT using cryomoulds (Tissue-Tek), frozen in dry ice cooled isopentane, and sectioned at 14 µm on a cryostat (Leica, CM3050S) onto Superfrost slides.

#### Chicken embryos

the embryos were scooped out of the egg and dissected in cold PBS. The electroporated section of the trunk was isolated under the fluorescence scope and then fixed in 4% PFA overnight. After PBS washes, specimens were cryoprotected in 15% sucrose and then 30% sucrose in PBS, until they sank. They were then embedded in OCT using cryomoulds (Tissue-Tek), frozen in dry-ice cooled isopentane, and sectioned at 7 µm on a cryostat (Leica, CM3050S) onto Superfrost slides.

### Immunohistochemistry and TUNEL staining

#### Mouse brains

Sections were incubator in PBS for 10 min and then blocked with blocking buffer (5% Bovine Serum Albumin (BSA, Sigma) in PBS with 0.1% Triton X-100). Primary antibodies (MEIS2, 1:3000) in blocking buffer were placed on slides overnight at 4^°^C, washed in PBS with 0.1% Triton X-100 (PBST) and applied with secondary antibodies (1:500 Alexa Fluor-conjugated secondary antibodies in blocking buffer) for 1 hour at room temperature. Counterstaining was performed using Hoechst 33258 (Invitrogen). The slides were then washed in PBST, then mounted with a coverslip and Fluorogel mounting medium (Electron Microscopy Sciences). TUNEL staining was performed prior to secondary antibody incubation using the Biotin-based in situ cell death detection kit (Roche) and following the manufacturer instructions, including an Avidin-Biotin blocking step (Vector Laboratories).

### In situ hybridization

The DIG-labeled RNA probe for *Ctgf* was prepared by *in vitro* transcription of a PCR-amplified template from embryonic day 12.5 cDNA, using the following primers: Fwd: 5’-CAGCTGGGAGAACTGTGTAC-3’, Rev(T7): 5’-CGATGTTAATACGACTCACTATAGGGGCTCGCATCATAGTTGGGTC-3’. Hybridization procedure as described in (Rosello-Diez et al., 2018).

### Skeletal preparations

After embryo collection, the skin, internal organs and adipose tissue were removed. The samples were then fixed in 95 % EtOH overnight at room temperature. To remove excess fat, the samples were then incubated in acetone overnight at room temperature. To stain the cartilage, the samples were submerged in a glass scintillation vial containing Alcian blue solution (0.04 % (w/v), 70 % EtOH, 20 % acetic acid) and incubated at least overnight at room temperature. The samples were destained by incubating them in 95 % EtOH overnight, and then equilibrated in 70% EtOH, prior to being pre-cleared in 1 % KOH solution for 1-10h at room temperature (until blue skeletal elements were seen through). The KOH solution was replaced with Alizarin red solution (0.005 % (w/v) in water) for 3–4h at room temperature. The Alizarin red solution was then replaced with 1-2% KOH until most soft tissues were cleared. For final clearing, the samples were progressively equilibrated through 20% glycerol:80% (1%KOH), 50 % glycerol:50 % (1 %) KOH and finally transferred to100 % glycerol for long-term storage.

### Image Analysis and Quantification

The regions of interest were isolated from imaged sections of the multiple models. All the parameters including the brightness, contrast, filters, layers were kept the same for all the images in the same study. Cells positive for the signal of interest were counted manually using ImageJ, after acquisition on an Imager Z1 microscope (Zeiss).

### Statistical Analysis

The comparisons were done by an unpaired Mann-Whitney test (1 variable and 2 conditions), or by 2-way ANOVA (2 variables and 2 or more conditions) following a matched (paired) design when possible (indicated when not). When left and right measurements were compared within experimental animals only, Wilcoxon matched-pairs signed rank test was used. Non-parametric tests were chosen when normality could not be confirmed. All the statistical analyses were done using Prism7 software (Graphpad). No blinding was used except for Fig. 6C. Electroporated chicken embryos were randomly assigned to control or experimental groups. No previous power analysis was done to estimate the sample size.

## ACKNOWLEDGEMENTS

We thank members of the Joyner and Rosello-Diez laboratories for discussions and technical help. We are grateful to Dr. Miguel Torres for providing a polyclonal antibody to all isoforms of mouse MEIS2, and to Megan Davey for sharing her expertise with TAT-Cre beads. We acknowledge the assistance from Monash Animal Research Platform (MARP), Monash Micro Imaging Platform (MMI) and the Sloan Kettering Institute Research Animal Resource Center and Mouse Genetics Core Facility. We also thank the Center for Comparative Medicine and Pathology and the Mouse Genetics Core Facilities of MSKCC for outstanding technical support. Plasmid *pT2-CAGGS-H2B-BFP* was a gift of the Marcelle lab. Addgene plasmids #20910 (*PB-CA-rtTA* Adv) and #102423 (*PB-CAGGS-rtTA-IRES-Hygro*) were donated by the Nagy and Koh labs, respectively.

## AUTHOR CONTRIBUTIONS

E.A.: performed some *Tigre*^*Dragon-Ctgf*^ and *Tigre*^*Dragon-DTA*^ experiments and most chicken experiments and edited the manuscript; A.S.: performed data analysis and edited the manuscript; X.Q.: assisted with genotyping, cloning, embryo dissection, histology and data acquisition; N.S.B.: performed cerebellar experiments and analysis, D.S. assisted with breeding, genotyping and histology throughout the project; L.M. and H.Z.: performed ES cell targeting and provided key reagents; A.L.J was involved in experimental design, funding acquisition, supervision and manuscript editing; A.R-D. was responsible of conceptualization, funding acquisition, experimental design and procedures, data acquisition and analysis, supervision, manuscript drafting and editing.

