## Supplementary for "A collection of genetic mouse lines and related tools for inducible and reversible intersectional misexpression"

### SUPPLEMENTARY FIGURES

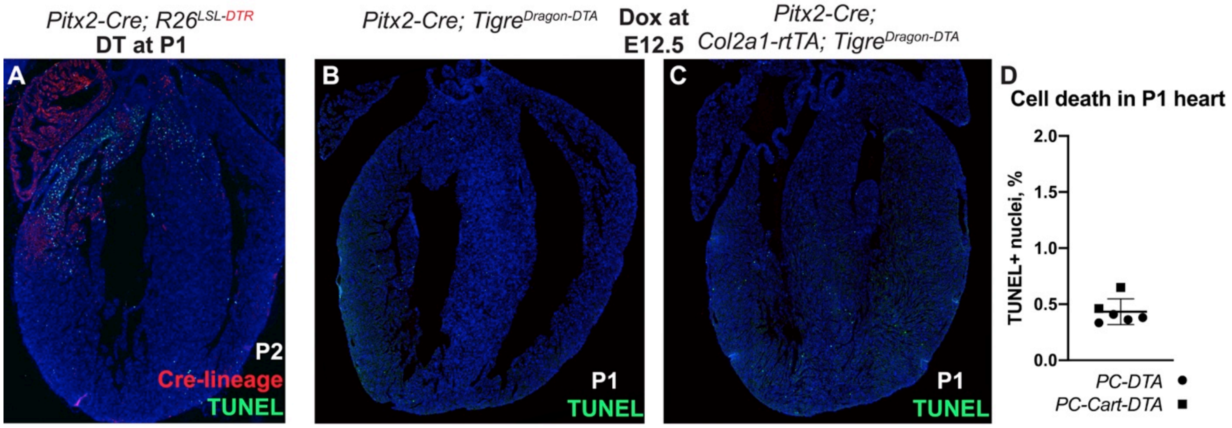


**Supplementary Figure 1. The intersectional approach allows sparing of unwanted cell populations, such as cardiomyocytes when using *Pitx2-Cre*.** A) Representative image showing the cell death distribution (TUNEL) in the P2 heart when diphtheria toxin receptor (DTR, red) is activated by *Pitx2-Cre* and DT is injected at P1 (modified from (Rosello-Diez et al., 2017), subject to a free-distribution license). B, C) Negligible cell death is observed in the hearts of P1 *PC-DTA* (B) and *PC-Cart-DTA* (C), with Dox provided at E12.5. D) Quantification of cell death in *PC-DTA* and *PC-Cart-DTA* hearts (mean±SD, n=4 and 2 respectively).

### SUPPLEMENTARY MATERIALS AND METHODS

| REAGENT or RESOURCE | SOURCE | IDENTIFIER |
|---|---|---|
| **Antibodies** | | |
| rat monoclonal anti-mCherry | ThermoFisher | M11217 mAb; RRID:AB_2536611 |
| rabbit polyclonal anti-p21 | Santa Cruz | M19 pAb; RRID:AB_632123 |
| Rabbit polyclonal anti-MEIS2 | Mercader et al., 2005 | N/A |
| Anti-Digoxigenin-AP Fab fragments | Sigma-Aldrich | 11093274910; RRID:AB_514497 |
| **Chemicals, recombinant proteins** | | |
| Paraformaldehyde (powder) | Sigma-Aldrich | |
| Triton X-100 | Sigma-Aldrich | T9284-500ML |

| | | |
|---|---|---|
| Fluoromount aqueous mounting medium | Sigma-Aldrich | F4680-25ml |
| 40,6-diamidino-2-phenylindole (DAPI) | ThermoFisher | D1306 |
| SYTOX green | ThermoFisher | S7020 |
| Doxycycline hyclate | Sigma-Aldrich | D9891-10 gr |
| Affigel blue beads | Biorad | 1537301 |
| Biotin 16-dUTP | Sapphire Bioscience | JBS-NU-803-BIO16-S |
| TAT-Cre | Millipore | SCR-508 |
| Terminal transferase recombinant | Merck | 3333566001 |
| **Electroporation tools** | | |
| Platinum wire, 0.25 mm | SDR Scientific | 711000 |
| Tungsten wire, 0.5 mm | Sigma-Aldrich | 356972 |
| Pulse generator and amplifier | Intracel | TSS20 |
| Glass capillaries | Harvard apparatus | GC120T-10 |
| **Experimental Models: Organisms/Strains** | | |
| Mouse $Tigre^{Dragon\text{-}p21}$ | Rosello-Diez et al., 2018 | RRID:IMSR_JAX:034777 |
| Mouse $Tigre^{TRE\text{-}p21}$ | This paper | N/A |
| Mouse $Tigre^{Dragon\text{-}DTA}$ | This paper | RRID:IMSR_JAX:034778 |
| Mouse $Tigre^{Dragon\text{-}Ctgf}$ | This paper | RRID:IMSR_JAX:034779 |
| Mouse B6.Cg-Gt(ROSA)26Sortm1.1 (CAG-rtTA3)Slowe/LdowJ | Dow et al., 2014 | RRID:IMSR_JAX:029627 |
| Mouse *Atoh1-tTA* | Willett et al., 2019 | N/A |
| Mouse *TetO-Cre* | Perl et al., 2002 | RRID:IMSR_JAX:006234 |
| Mouse *En1-Cre* | Kimmel et al., 2000 | RRID:IMSR_JAX:007916 |
| Mouse *Col2a1-tTA* | Rosello-Diez et al., 2018 | N/A |
| B6C3(129)-Tg(Pitx2ASE-cre)16Hmd/HmdRbrc Mus musculus | Shiratori et al., 2006 | RRID:IMSR_RBRC03487 |
| Tg(Col2a1-rtTA)1Jath | Posey et al., 2009 | RRID:MGI:4361029 |
| **Plasmids** | | |
| pDragon-p21 | Rosello-Diez et al., 2018 | RRID:Addgene_140894 |
| pDragon-DTA | This paper | RRID:Addgene_140895 |
| pDragon-Ctgf | This paper | RRID:Addgene_140896 |
| pPB-CA-rtTA-Adv | Addgene | RRID:Addgene_20910 |
| pPB-CAG-rtTA-IRES-Hygro | Addgene | RRID:Addgene_102423 |
| pT2-CAG-H2B-TagBFP | Sieiro et al., 2016 | N/A |
| pCAGGs-rtTA-IRES-Cre | This paper | AR98 |
| Ai62(TITL-tdT) Flp-in replacement vector | Madisen et al., 2015 | RRID:Addgene_61576 |

**Supplementary Table 1. List of reagents and sources**

| Allele | Program | Primer name | Primer sequence (5' to 3') |
|---|---|---|---|
| Cre | 94ºC 2', 33x(94ºC 20", 60ºC 20", 72ºC 30"), 72ºC 5', 10ºC forever) | Cre F<br>Cre R | GCGGTCTGGCAGTAAAAACTATC<br>GTGAAACAGCATTGCTGTCACTT |
| Dragon WT | 94ºC 2', 33x(94ºC 20", 55ºC 20", 72ºC 30"), 72ºC 5', 10ºC forever) | Dragon F<br>Dragon R WT | CCCAACGGTCACTTACTTCC<br>CACACCTTTAATCCCGATGC |
| Dragon Mut | 94ºC 2', 10x(94ºC 20", 65ºC-0.5/cycle 15", 68ºC 10"), 72ºC 5', 10ºC forever), 28x(94ºC 15", 60ºC 15", 72ºC 10"), 72ºC 2', 10ºC forever) | Dragon F<br>Dragon R Mut | CCCAACGGTCACTTACTTCC<br>GGTAACCGCGGCATAAAAC |
| p21rec | 94ºC 2', 33x(94ºC 20", 55ºC 20", 72ºC 30"), 72ºC 5', 10ºC forever) | Pmin F<br>WPRE Rev | AGTGAACCGTCAGATCGC<br>GCGTATCCACATAGCGTAAAAG |
| Col2-rtTA | 94ºC 2', 33x(94ºC 20", 61ºC 20", 72ºC 30"), 72ºC 5', 10ºC forever), | rtTA F<br>rtTA R | ATGCCCTTGGAATTGACGAGTACGG<br>CGAGGCTTGCAGGATCATAATCAG |
| Col2-tTA | 94ºC 2', 33x(94ºC 20", 55ºC 20", 72ºC 30"), 72ºC 5', 10ºC forever) | Col2a1 F<br>tTA R | CCAGGGTTTCCTTGATGATG<br>GCTACTTGATGCTCCTGATCCTCC |
| CAG-rtTA3 | 94ºC 2', 33x(94ºC 20", 55ºC 20", 72ºC 30"), 72ºC 5', 10ºC forever) | Mut F<br>Common Rev<br>WT F | AAAGTCGCTCTGAGTTGTTAT<br>GCGAAGAGTTTGTCCTCAACC<br>GGAGCGGGAGAAATGGATATG |

**Supplementary Table 2. Primers and PCR conditions used for genotyping**
